# A Novel VWF Knockout Endothelial Cell Model to study von Willebrand Factor Biology and Von Willebrand Disease Mechanisms

**DOI:** 10.64898/2026.04.01.715845

**Authors:** Isabel Bär, Petra E. Bürgisser, Buram Ardic, Jeroen C.J. Eikenboom, Jan Voorberg, Frank W.G. Leebeek, Ruben Bierings

**Affiliations:** Hematology, Erasmus University Medical Center, Rotterdam, The Netherlands; Internal Medicine, Division of Thrombosis and Hemostasis, Leiden University Medical Center, Leiden, The Netherlands; Molecular Hematology, Sanquin Research, Amsterdam, The Netherlands; Experimental Vascular Medicine, Amsterdam University Medical Center, Amsterdam, The Netherlands

## Abstract

Understanding how specific VWF variants disrupt endothelial processing and function is central to elucidating von Willebrand disease (VWD) pathophysiology. However, current in vitro systems lack either the endothelial specificity or the genetic flexibility required for systematic variant characterization. Here, we present a genetically defined VWF-knockout cord-blood–derived endothelial colony-forming cell (VWF-KO cbECFC) model that enables controlled reintroduction of VWF variants in a physiologically relevant endothelial context. Using a patient with type 3 VWD carrying the homozygous pathogenic variant p.M771V and a second homozygous variant of uncertain significance p.R2663P as a reference, we demonstrate that expression of p.M771V in VWF-KO cbECFCs reproduces the patient’s intracellular processing defect and loss of high-molecular-weight multimers, whereas p.R2663P behaves as a benign allele. These findings establish the model’s ability to accurately distinguish pathogenic from non-pathogenic variants. Comparative analyses with HEK293 cells show that VWF-KO cbECFCs provide superior subcellular resolution, reliably forming authentic Weibel–Palade bodies (WPBs) and faithfully revealing ER retention phenotypes that remain ambiguous in non-endothelial systems. The proliferative capacity of cbECFCs further enables scalable and reproducible experimentation, overcoming major limitations associated with patient-derived ECFCs.

Looking ahead, the VWF-KO cbECFC platform offers broad potential for VWF and VWD research. Its endothelial identity and genetic flexibility make it suitable for investigating VWF biosynthesis and trafficking, secretion dynamics, WPB biology, angiogenic processes, and shear-dependent VWF function. This system therefore provides a versatile foundation for mechanistic studies, systematic variant assessment, and future translational applications.

## Introduction

Von Willebrand disease (VWD) is the most common inherited bleeding disorder, affecting approximately 1% of the global population^1,2^. It results from quantitative or qualitative defects in von Willebrand factor (VWF), a large glycosylated multimeric protein essential for primary and secondary hemostasis^3^. VWF mediates platelet adhesion at sites of vascular injury and serves as a carrier for coagulation factor VIII^4^. The functional integrity of VWF depends on its assembly into high molecular weight (HMW) multimers, which are stored in endothelial-specific organelles known as Weibel-Palade bodies (WPBs) that can be secreted upon endothelial activation^5,6^. Formation of WPBs is dependent on VWF synthesis as illustrated by their loss in VWF-deficient endothelial cells^7–11^ and the formation of so-called pseudo-WPBs after ectopic expression in non-endothelial cell types^12–14^. Disruption of VWF synthesis, multimerization, or secretion leads to a heterogeneous spectrum of bleeding phenotypes and is often accompanied by abnormal WPB biosynthesis or morphology in endothelial cells^15^. To date, more than 750 pathogenic VWF variants have been identified^16^, yet many remain poorly characterized and rely on predictive algorithms for pathogenicity assessment. Understanding the molecular mechanisms underlying these variants is necessary for improving diagnosis and enabling personalized treatment strategies^17^.

Several cellular models have been employed to study VWF biology and VWD. HEK293 cells are widely used because they lack endogenous VWF, allowing controlled ectopic expression of VWF variants. While this system allows for analysis of VWF post-translational processing and multimer formation, it poorly reproduces key endothelial features such as regulated secretion. Moreover, the pseudo-WPBs in HEK293 cells often cluster in the perinuclear area where the organelles can be difficult to visually separate, which complicates comparison of their morphology to that of their authentic endothelial counterparts. Consequently, despite their long-standing use as a model system for VWF and VWD biology, HEK293 cells fall short when it comes to studying intracellular trafficking and secretion defects associated with VWD variants. Endothelial colony-forming cells (ECFCs) derived from venous blood of VWD patients represent a more physiologically relevant model, as they carry the patient’s genetic background and exhibit authentic endothelial characteristics^18^. However, patient-derived ECFCs can be difficult to obtain and have limited proliferative capacity, restricting their use to a few passages and complicating large-scale or long-term studies^19^. Cord blood-derived ECFCs (cbECFCs) offer enhanced proliferative capacity compared to peripheral blood ECFCs, however, obtaining umbilical cord blood from a patient is even more challenging due to the limited availability.

To overcome these limitations, we developed a novel endothelial cell model based on cbECFCs, where endogenous VWF expression is knocked out using CRISPR/Cas9. This system provides a stable platform for introducing VWF variants into a *bona fide* endothelial cell model so that phenotypes can be studied in their native context without interference of endogenous VWF expression. In this study, we characterize the VWF KO cbECFC model and demonstrate its ability to replicate disease-specific phenotypes. Specifically, we investigate two variants (p.M771V and p.R2663P) that were previously identified in a patient with severe VWD3^20^, and determine how their phenotypes compare when expressed in VWF KO cbECFCs and HEK293 cells. The resulting phenotypes were also evaluated against patient-derived ECFCs to assess how closely each model reflects the authentic endothelial phenotype. This approach allowed us to validate the utility of VWF KO cbECFCs as a reliable and versatile tool for mechanistic studies of VWF biology and VWD pathogenesis.

## Methods

### Patients

Patients were included in the WiNpro study (Willebrand in the Netherlands study), which is a nationwide, prospective cohort study in patients with VWD (ClinicalTrials.gov: NCT03521583). The inclusion and exclusion criteria for the WiNpro study are further explained in the supplemental materials. One severe VWD patient carrying both the p.M771V and p.R2663P variants and a healthy umbilical-cord donor gave informed consent to isolate and study their ECFCs in the 2020-BOEC-MK (NL72564.078.20; MEC-2020-0214) and the 2019-UMBILICAL-CORD (MEC-2019-0699) studies, which were performed in accordance with the Declaration of Helsinki.

### ECFC isolation and culturing of cells

ECFCs and cord-blood ECFCs were isolated from venous blood or umbilical cord blood, respectively, as previously described^21^. The ECFCs were cultured on 1% gelatin coated flasks in EGM-2 (Promocell, C22011) containing 18% FCS (Bodino) and 5% penicillin/streptomycin.

LentiX-HEK293T cells (Takara, Cat. No. 632180) were used for virus production and cultured in Dulbecco modified Eagle medium containing D-glucose and L-glutamine (Thermo Fisher Scientific, 11965092), supplemented with 10% FCS and 5% penicillin/streptomycin. All cells were cultured at 37°C and 5% CO2 in a humidified atmosphere and passaged using 0.05% Trypsin-EDTA.

### Plasmid constructs and viral delivery

A CRISPR/Cas9-v2 construct containing a Blasticidin S cassette and a SpCas9 (Addgene #83480^22^) was used in this study to introduce double strand breaks (DSBs), leading to the introduction of indels that disrupt the VWF open reading frame and a gene knock-out. Complementary oligos with BsmBI-compatible overhangs were ordered from Integrated DNA Technologies (IDT), encoding the gRNA sequence 5’-AGCACCCCGGCAAATCTGGC-3’ targeting the second exon of VWF (Figure 1A).

**Figure 1:**
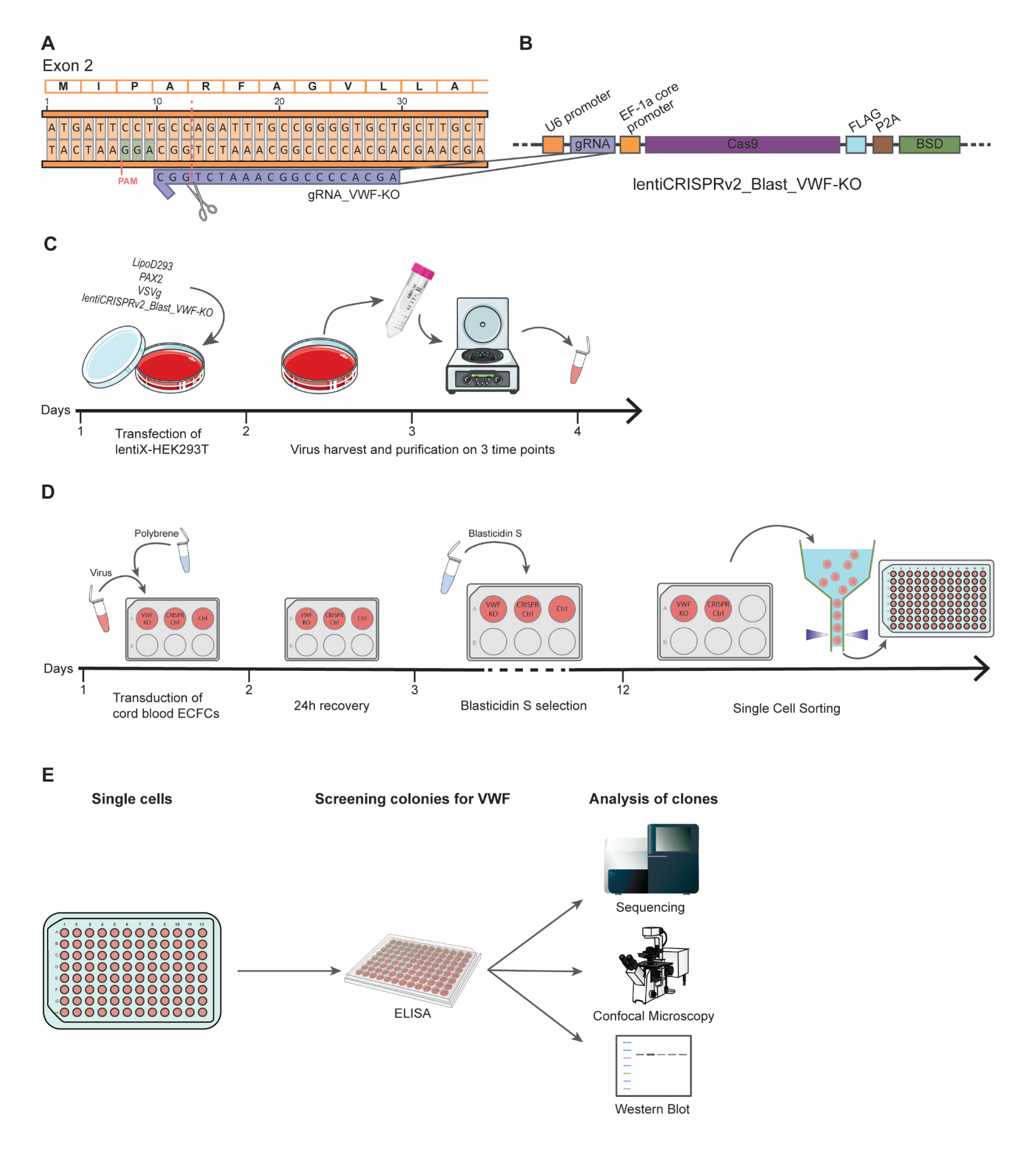
Workflow of generating VWF KO cell lines from a healthy cord blood ECFC clone. A) gRNA location on Exon 2 of the *VWF* gene. Predicted CRISPR/Cas9 cutting site at c.G12 is indicated. B) Plasmid map of lentiCRISPRv2 vector carrying a Blasticidin S resistance. C) Workflow of virus generation in lentiX-HEK293T cells and virus purification. D) Workflow of knocking out VWF in cbECFCs. E) Screening and validation of VWF knock-out cell lines generated from a single cell.

The wild-type VWF expression construct was based on the pcDNA3.1-VWF-WT backbone. To introduce the p.M771V and p.R2663P variants, we ordered synthetic DNA fragments (Base-Gene) containing the respective mutations. These fragments were cloned into the pcDNA3.1-VWF-WT vector using standard restriction enzyme digestion and ligation techniques, followed by transformation into competent *Stbl3 E. coli* cells for amplification. Plasmid integrity and correct insertion of the mutations were confirmed by Sanger sequencing.

For the production of lentiviral particles, lentiX-HEK293T cells were transfected with LipoD293 (Ver.II, SignaGen) according to the instructions of the supplier. A generation II packaging system with pVSVg and psPAX2 (Addgene #12259 and #12260, gift from Professor Didier Trono) was used. Harvesting was performed 24h, 48h, and 72h after transfection. The virus harvests were concentrated in Amicon Filter Units (Merck) and stored at -80 °C until use (Figure 1C).

Transduction of the cells was performed in the presence of 0.4 µg/ml polybrene (Merck). Enrichment was achieved using 5 ug/ml blasticidin S for 10 days for the VWF-KO in cbECFCs, and 0.6 µg/ml puromycin (Merck) for 4 days for the overexpression in the VWF-KO cbECFC cell line.

### DNA isolation and Sanger Sequencing

Genomic DNA was isolated using DNA lysisbuffer (100mM Tris, 0.2% SDS, 6mM EDTA and 200mM NaCl) and a subsequent incubation of at least 1h at 55°C shaking. Precipitation with isopropanol was performed at a 1:1 ratio. After centrifugation at 13 000g for 10 mins the DNA pellet was washed twice with 80% EtOH and eluted in TE buffer (Invitrogen). The DNA was analyzed by Sanger sequencing at Macrogen Europe.

### Immunocytochemistry and image analysis

Cells were fixed and stained as previously described^20^. The antibodies and dilutions are listed in **Table 1**.

**Table 1:** Antibodies used in this study.

| Primary<br>Antibody | Supplier; Lot<br>Nr. | Dilution | Secondary<br>Antibody | Supplier; Cat.<br>Nr. |
| --- | --- | --- | --- | --- |
| **Polyclonal**<br>**rabbit anti**<br>**human VWF** | DAKO/Agilent,<br>LOT0083511 | 1:50 000 | Donkey anti<br>Rabbit IgG | Biotium; 20015 |
| **Monoclonal**<br>**mouse anti**<br>**human PDI** | Sigma/Merck;<br>LOT<br>MAB03333 | 1:1000 | Goat anti<br>Mouse IgG1 | Biotium; 20248 |
| **polyclonal**<br>**goat anti**<br>**human VE-**<br>**cadherin** | R&D systems;<br>LOT<br>EF0321071 | 1:300 | Donkey anti<br>Goat IgG | Thermo<br>Fisher;<br>A21447 |

### Western Blotting

Single-cell sorted clones of VWF-KO cbECFCs were lysed in NP-40 buffer (5 mM EDTA pH 8.0, 10 mM Tris-HCl pH 6.7, 150 mM NaCl, 0.5% Igepal) supplemented with protease inhibitor (Sigma, S8820). Protein concentrations of lysate samples were quantified using the Pierce 660 nm protein assay. Equal protein concentrations of lysate samples were separated on a gradient Bis-Tris NuPAGE gels (Invitrogen, 4% to 12%) under reducing conditions and transferred to a 0.2 μm nitrocellulose membrane using wet-blotting. Protein detection was facilitated using rabbit polyclonal anti-VWF and antibodies. Membranes were visualized on an Odyssey Clx scanner (Li-COR).

### Multimer analysis

Conditioned media of ECFCs were analyzed for their VWF multimer pattern as described previously^23^. Briefly, samples in Tris/EDTA and glycerol were heated to 60⁰C for 20 minutes before running them on a 0.9% agarose gel at 75V for 6 hours at 4⁰C. The gel was then reduced in β-mercaptoethanol in PBS for 10 minutes on a shaker and subsequently washed with PBS. The gel was transferred to a 0.2 µm nitrocellulose membrane by capillary suction overnight. After blocking in 5% BSA, multimer patterns were visualized using VWF-HRP AB and ECL solution (Thermo Fisher Scientific SuperSignalTM West Pico PLUS Chemiluminescent Substrate) on an Amersham Imager 600. ImageJ64 was used to translate the multimer pattern into densitometry graphics^24^.

## Results

### Generation of a novel endothelial cell VWF knock-out model

To establish a cellular model for von Willebrand disease (VWD) research, we generated VWF-deficient endothelial cells by knocking out VWF in cord blood-derived endothelial colony-forming cells (cbECFCs). Previous studies have demonstrated that genetic modification of ECFCs using CRISPR/Cas9 does not compromise their endothelial characteristics^11,20,25^. Building on this, we performed CRISPR/Cas9-mediated VWF knockout in long-lived cbECFCs.

After evaluating 3 gRNAs targeting exon 1, exon 2, and exon 3, we selected the guide RNA targeting exon 2 of VWF to introduce a double-strand break at the 12th transcribed base of the gene (Figure 1A). The CRISPR/Cas9 vector included a Blasticidin S resistance cassette to enable selective enrichment of edited cells while avoiding Puromycin resistance, which is commonly used in overexpression systems (Figure 1B). The transduction was performed with purified lentivirus (Figure 1C). The choice of cbECFCs over venous blood-derived ECFCs was critical, as the knockout procedure including single-cell sorting requires robust proliferative capacity, which is limited with venous blood-derived ECFCs (Figure 1D). Candidate VWF-KO clones were initially screened using a VWF ELISA on conditioned media (data not shown) and subsequently validated by Sanger sequencing, confocal microscopy, and Western Blot (Figure 1E).

Out of 10 tested clones, 9 showed a loss of VWF protein expression in lysates, demonstrating a successful VWF KO (Figure 2A). We selected clone 10 for downstream experiments. Phenotypically, confocal microscopy showed absence of VWF staining in the cell periphery, while VE-cadherin remained detectable at the cell-cell junctions, confirming preservation of endothelial identity (Figure 2B). Analysis of conditioned medium from Clone 10 using a multimer assay (Figure 2C) and densitometry (Figure 2D) revealed loss of HMW VWF multimers. The LMW VWF oligomers that were detected originated from the high fetal calf serum content in the endothelial growth medium (EGM-18) and were therefore also present in the medium only sample. Genetic disruption of VWF was confirmed by Sanger sequencing, which revealed deletion within exon 2 (Figure 2E). The Sanger sequencing for the remaining 8 clones showing no protein expression in Western Blot is shown in supplemental figure 1.

**Figure 2:**
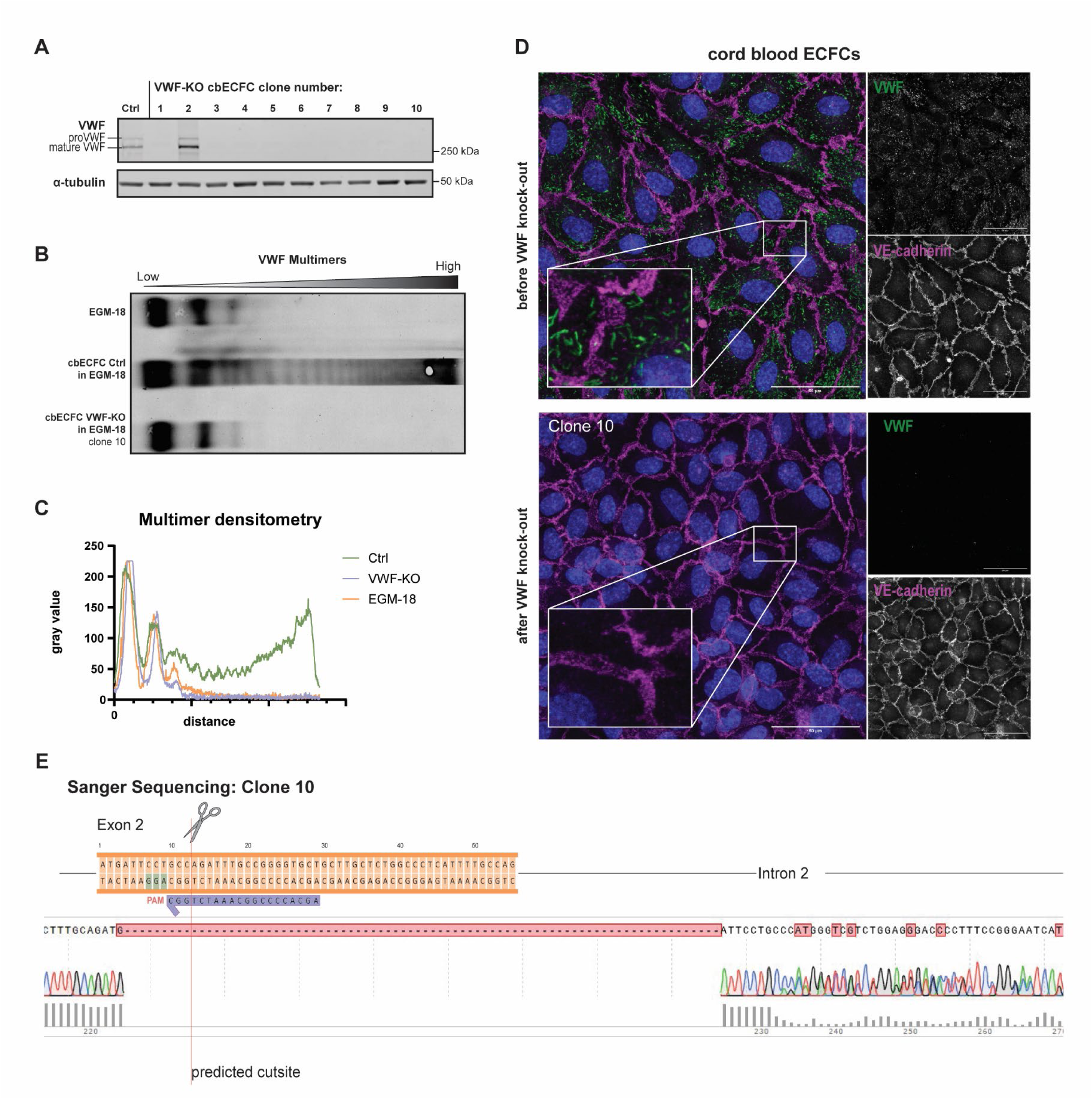
Confirmation of successful VWF knock-out in single cell derived clones of cord blood ECFCs. A) Western Blot analysis of 10 cbECFC clone lysates after CRISPR/Cas9 treatment and single cell sorting, revealing successful VWF protein loss in 9 out of 10 clones. B) Multimer analysis of secreted VWF into supernatant of untreated cbECFCs and clone 10. A medium control (EGM-18) is loaded to show that low molecular weight VWF is present in the serum of the growth medium and therefore always present in secretome samples. C) Multimer densitometry of the multimer analysis depicted in B. D) Confocal analysis of cbECFCs before CRISPR/Cas9 treatment and of clone 10 after VWF KO. Blue: DAPI, Green: VWF, Magenta: VE-cadherin. Scalebar: 25µm E) Genetic analysis of clone 10 using Sanger sequencing to prove successful CRISPR/Cas9 cutting and shift of the ORF.

### VWF-KO cbECFCs accurately model patient-specific biochemical defects in VWF processing upon variant introduction

To establish a reference phenotype for variant interpretation, we used a previously characterized VWD patient carrying the pathogenic homozygous VWF p.M771V variant, associated with a severe bleeding phenotype^20^. Clinically, the patient was diagnosed with severe von Willebrand disease, presenting with a high ISTH-BAT score of 33 (Figure 3A). In addition to the homozygous pathogenic p.M771V variant, sequencing revealed a second homozygous variant, p.R2663P, of which the clinical significance remains uncertain according to conflicting classifications in common databases^26^. In our earlier work, we connected the strong bleeding phenotype with a intracellular processing defect of VWF. The p.M771V variant is located in proximity of the furin cleavage site and the insufficient cleavage of VWFpp from the proVWF variant prevents further processing into mature VWF and therefore a complete loss of HMW VWF^20^. To evaluate whether the newly generated VWF-KO cbECFC line can reproduce these variant-specific abnormalities, we reintroduced VWF WT (control), p.M771V (known pathogenic variant), or p.R2663P (variant of unknown significance) into the knockout background. Since the multimeric composition of secreted VWF reflects the efficiency of intracellular processing and the formation of HMW multimers stored in WPBs, we first assessed VWF secretion of the cbECFCs by multimer analysis. Expression of VWF WT led to a complete VWF multimer ladder, whereas expression of p.M771V led to a loss of HMW multimers (Figure 3B), identical to the defect observed in patient-derived p.M771V ECFCs. In contrast, VWF-KO cbECFCs expressing p.R2663P preserved the full VWF multimer ladder, indicating that this variant does not impair VWF biosynthesis or multimerization. At the intracellular level, the p.M771V mutation caused a shift from mature VWF to proVWF, reflecting the underlying processing defect. This was evident in the lysate immunoblots through the absence of a mature VWF band in VWF-KO cbECFCs expressing p.M771V, mirroring the absence observed in the two patient-derived ECFC lysate samples (Figure 3C-E). In contrast, VWF-KO cbECFCs expressing VWF WT or VWF p.R2663P showed two distinct bands, corresponding to both proVWF and mature VWF, consistent with normal intracellular processing. Together, these extracellular and intracellular biochemical findings demonstrate that VWF-KO cbECFCs faithfully reproduce patient-specific processing defects and provide a robust system for distinguishing pathogenic from benign VWF variants.

**Figure 3:**
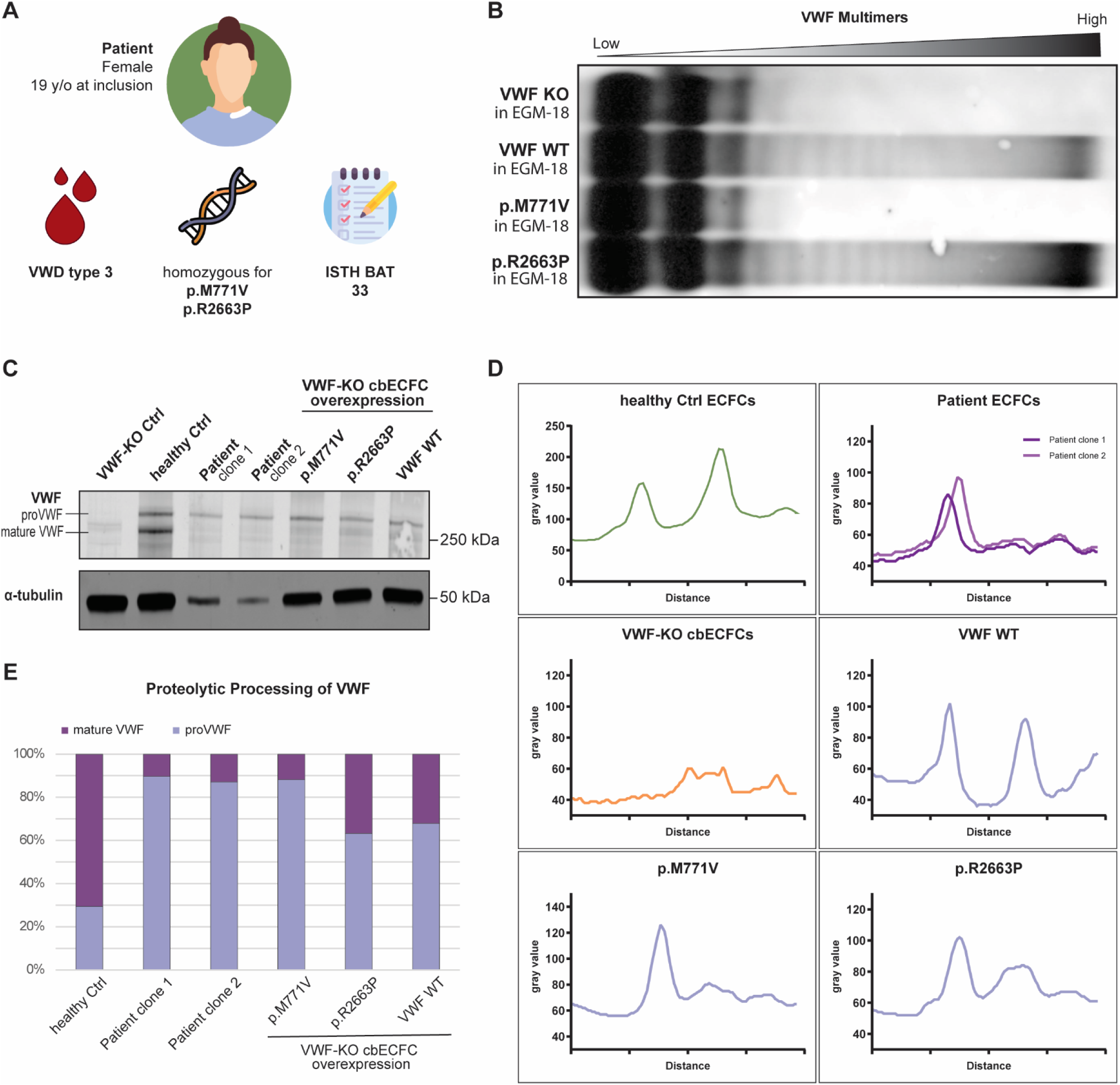
VWF-KO cbECFCs accurately model patient-specific VWF processing defects. A) Characteristics of a female VWD type 3 patient B) Multimer analysis of secreted VWF by the VWF-KO cbECFCs and the reinstated VWF WT, VWF p.M771V, and VWF p.R2663P conditions. LMW multimers are artifacts from the EGM-18 medium and not corresponding to secreted VWF by the cells. C) Western blot of lysates from ECFCs. The two patient samples represent two different ECFC colonies of the same patient. D-E) Densitometry analysis of the western blot depicted in C. The first peak corresponds to pro VWF, and the second peak corresponds to mature VWF.

### VWF-KO cbECFCs generate authentic WPBs and outperform HEK293 cells in mimicking VWF variant patient phenotypes

To evaluate the suitability of VWF-KO cbECFCs as a cellular system for analyzing VWF variant behaviour, we compared intracellular VWF localization and storage in these endothelial cells with that observed in HEK293 cells expressing the same constructs (Figure 4). Both conditions were compared to the prominent features of the patient-derived venous blood ECFCs: The patient ECFCs carrying the p.M771V and p.R2663P variants exhibited pronounced VWF retention within the endoplasmic reticulum (ER) and a complete absence of WPBs in the cells, which is consistent with the severe intracellular trafficking and processing defect we previously observed^20^. In HEK293 cells, VWF distribution appeared heterogeneous and often difficult to interpret. Under VWF WT conditions, some cells displayed slightly elongated structures reminiscent of WPBs (white arrows), whereas other areas in the same cell exhibited predominantly diffuse VWF staining without clear organelle identity (orange arrows) (Figure 4B). Similar variability was observed for the p.R2663P and p.M771V variant. Co-localization analysis showed low overlap between VWF WT and PDI (Pearson’s r ≈ 0.1), while p.M771V exhibited strong overlap (r ≈ 0.6); however, the compact intracellular architecture of HEK293 cells limited the interpretability of ER retention and could explain the uncertain co-localization in the p.R2663P condition (r ≈ 0.3).

**Figure 4:**
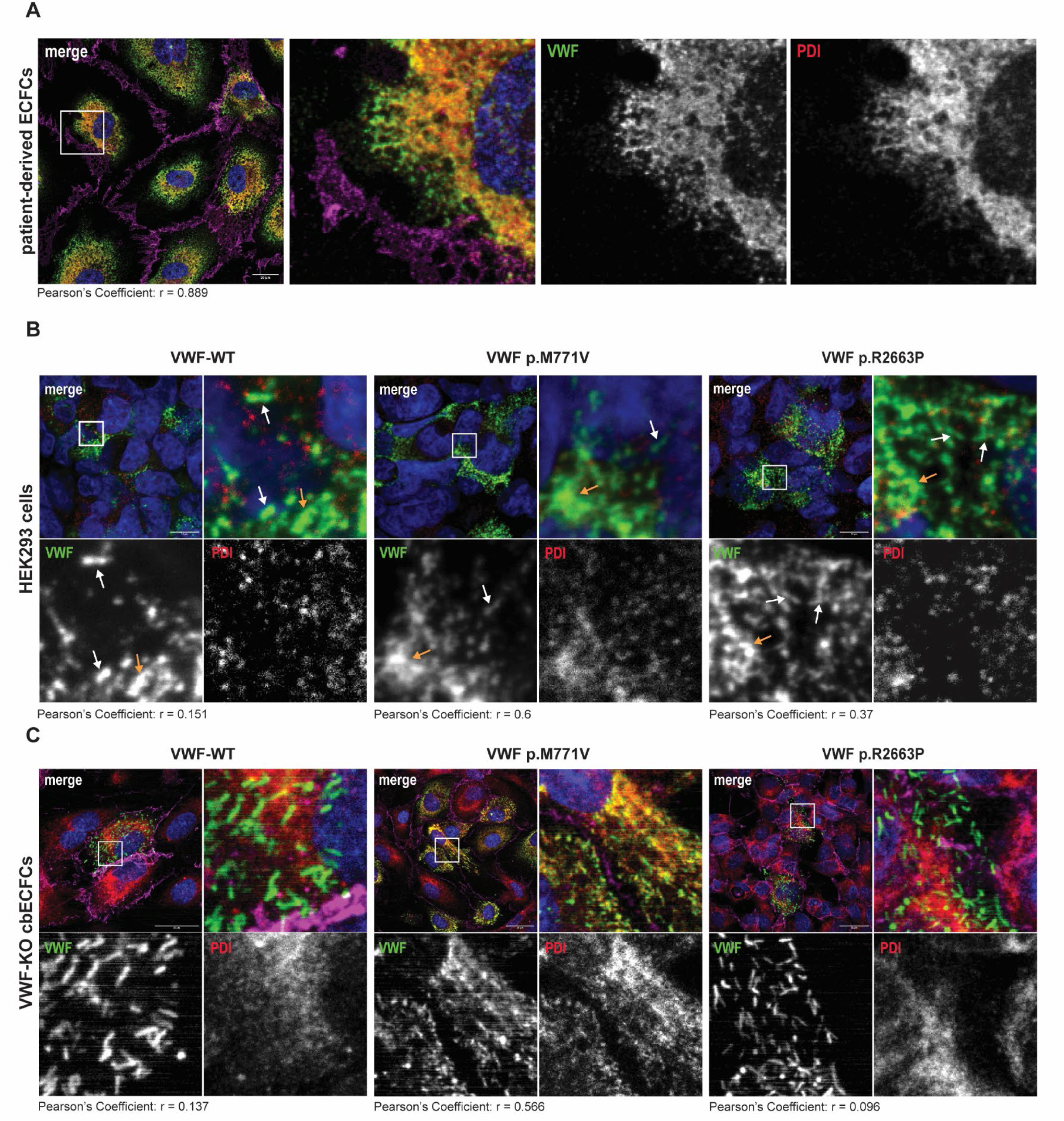
VWF-KO cbECFCs provide superior intracellular phenotypic resolution compared to HEK293 cells. A) Phenotype of patient-derived venous blood ECFCs showing a strong Pearson’s correlation between VWF and PDI of r=0.889. scalebar = 20µm B) Overexpression of VWF-WT, VWF p.M771V, and VWF p.R2663P in HEK293 cells. Pearson’s Correlation between VWF and PDI was low for VWF WT (r=0.151), high for p.M771V (r=0.6), and unclear in p.R2663P (r=0.37) condition. ER retention can therefore be confirmed in the p.M771V condition, but neither confirmed nor be denied for the p.R2663P condition. Scalebar = 10 µm C) Overexpression of VWF-WT, VWF p.M771V, and VWF p.R2663P in VWF-KO cbECFCs. Pearson’s correlation is low in VWF-WT (r=0.137), high in p.M771V (r=0.566) and low in VWF p.R2663P (r=0.096) conditions. ER retention is therefore only confirmed in the p.M771V condition. Scalebar = 25 µm Blue: DAPI, Green: VWF, Red: PDI, Magenta: VE-cadherin. Scalebar: 25µm

In contrast, VWF-KO cbECFCs provided a clear and physiologically relevant endothelial context for phenotypic VWF assessment. Reintroduction of VWF WT resulted in the formation of elongated, uniformly distributed WPBs characteristic of healthy endothelial cells (Figure 4C). Expression of p.M771V produced strong VWF retention within the ER and a complete absence of WPBs, with pronounced VWF–PDI co-localization (r ≈ 0.6). Cells expressing p.R2663P formed normal WPBs with minimal VWF–PDI co-localization (r < 0.1), allowing clear discrimination from the pathogenic variant.

Together, these observations demonstrate that VWF-KO cbECFCs offer superior subcellular resolution and phenotypic clarity compared to HEK293 cells. While HEK293 cells display diffuse and inconsistent VWF patterns that complicate interpretation and favor cherry-picking, VWF-KO cbECFCs faithfully reveal variant-specific effects on VWF processing, ER retention, and WPB biogenesis; providing a robust and physiologically relevant platform for intracellular VWF variant analysis.

## Discussion

The development of a VWF-KO cbECFC line provides an endothelial model that fills an important gap in VWD research by enabling VWF research within a physiological context that retains hallmark features of endothelial VWF biology. Although HEK293 cells are frequently employed for recombinant VWF studies, their non-endothelial background limits their suitability for mechanistic work^18^. In our hands, VWF expression in HEK293 cells produced heterogeneous intracellular localization patterns, ranging from diffuse staining to pseudo-WPB–like structures, complicating interpretation of VWF processing defects. This ambiguity became particularly apparent when analyzing the p.R2663P variant of uncertain significance^26,27^, where diffuse staining could be interpreted either as benign or as indicative of impaired processing. HEK293 cells do support multimer formation^28,29^ and therefore remain valuable for rapid screening or secretome-focused studies, but their inability to form bona fide WPBs and their lack of regulated secretion pathways restrict their utility for questions involving endothelial function.

For applications that require an endothelial context, such as functional studies of stimulated VWF secretion or angiogenesis, researchers have turned towards HUVECs. HUVECs are commercially available and exhibit a prolonged lifespan comparable to cbECFCs, which allows a knock-down of VWF using siRNAs to study processes related to low or absent VWF levels^30,31^. However, their undefined genetic background and polyclonality introduce variability, and while VWF knockdowns via siRNA are feasible, endogenous VWF cannot be completely controlled or replaced. Patient-derived venous blood or cord blood ECFCs remain the most physiologically relevant system, as they faithfully reflect the endogenous genotype and phenotype. While cbECFCs show an improved proliferation compared to venous blood ECFCs, cord blood is rarely available. Studies with cbECFCs therefore exist^32,33^, but are limited, and most ECFCs used in studies arise from venous blood. Yet, the ECFC’s limited availability, low colony-forming efficiency from venous blood^19^, restricted proliferative lifespan^34^, and clonal heterogeneity^35,36^ severely limit scalability for mechanistic studies or variant comparisons. We have previously used ECFCs from healthy donors to introduce patient missense variants directly into the endogenous VWF locus using a base-editor CRISPR/Cas9 system^20^. While this approach also results in a faithful representation of a VWF variant within an endothelial cell and even holds the advantage of endogenous promoter control, the application of a base editor is not generally suitable to mimic all VWF variants, and a homozygous genotype may be challenging to achieve.

The VWF-KO cbECFC system addresses these challenges by providing an endothelial cell platform that is both genetically defined and experimentally flexible. The knockout background allows precise introduction of VWF variants without interference from endogenous protein, while the proliferative capacity of cbECFCs offers a significantly wider experimental window than whole blood-derived ECFCs. The ability of these cells to recapitulate both intracellular processing defects and extracellular multimer abnormalities, as exemplified by the p.M771V variant, demonstrates that they maintain key aspects of endothelial VWF biology. This makes the model well suited for dissecting mechanisms of VWF biosynthesis, trafficking, secretion, WPB formation, and variant pathophysiology.

Its main limitation, relative to base-edited ECFCs, lies in the overexpression context, which may not fully reflect native VWF transcriptional regulation. However, together, these two models form a complementary toolkit: The VWF-KO cbECFCs optimized for mechanistic flexibility and variant throughput, the base-edited ECFCs for precise modeling of endogenous regulation.

Looking forward, the VWF-KO cbECFC system opens several avenues for future research that extend beyond the scope of this study. It provides a foundation for investigating fundamental mechanisms of VWF processing, including glycosylation^37^, and WPB biogenesis. The platform is also ideally suited for studying stimulated VWF secretion dynamics, such as histamine- or PMA-induced exocytosis, allowing analysis of secretion mode, quantity, and cargo specificity. Furthermore, because WPBs store multiple endothelial effector molecules, this system can be used to explore broader WPB-associated functions, including angiogenesis, inflammatory signaling, and leukocyte recruitment^5^. Because of their proliferative capacity, VWF-KO cbECFCs can be utilized for microfluidic experiments^38^. Under flow conditions, VWF-KO cbECFCs expressing defined variants could be used to model VWF string formation and platelet recruitment, enabling detailed characterization of variant-specific defects in shear-dependent function. Finally, this model provides a scalable system for comprehensive evaluation of VWD variants, offering the possibility to systematically test uncharacterized or newly identified mutations and to support translational applications, including preclinical gene-therapy development and drug discovery.

In summary, the VWF-KO cbECFC platform constitutes a robust and physiologically meaningful system for dissecting VWF biology and VWD pathogenesis. Its combination of genetic control, endothelial identity, and experimental versatility makes it a powerful complement to both classical heterologous systems and advanced genome-edited endothelial models, and positions it as a valuable tool for future mechanistic, functional, and translational studies.

## Acknowledgements

This work was supported by funding from The Netherlands Organization for Scientific Research, Domain Applied and Engineering Sciences (TTW/AES), “Connecting Innovators” Open Technology Program, project no. 18712. Work in the Bierings lab is supported by grants from the Landsteiner Stichting voor Bloedtransfusie Research (LSBR-1923, M.V.D.B.; LSBR-1707, R.B.). We thank Sophie Hordijk for help with ECFC isolation. We thank Iris van Moort for valuable feedback and general input throughout the project, and Calvin van Kwawegen for assistance with patient recruitment. We thank the patient and healthy controls for their contribution and participation in our study.

## Authorship Contributions

I.B. performed experiments, analyzed data and drafted the manuscript; P.E.B. and B.A. performed experiments and analyzed data; J.C.J.E., J.V. and F.W.G.L. designed the study and provided clinical input; R.B. designed the study, supervised research and drafted the manuscript. All authors have read and approved the final version of the manuscript.

## Supplementary Figure

Sanger sequencing analysis of the other clones depicted in Figure 2A with their genetic modifications at the CRISPR cutting site leading to a VWF-KO.

