## Supplementary figures and images for "A Novel VWF Knockout Endothelial Cell Model to study von Willebrand Factor Biology and Von Willebrand Disease Mechanisms"

### Supplementary Figure

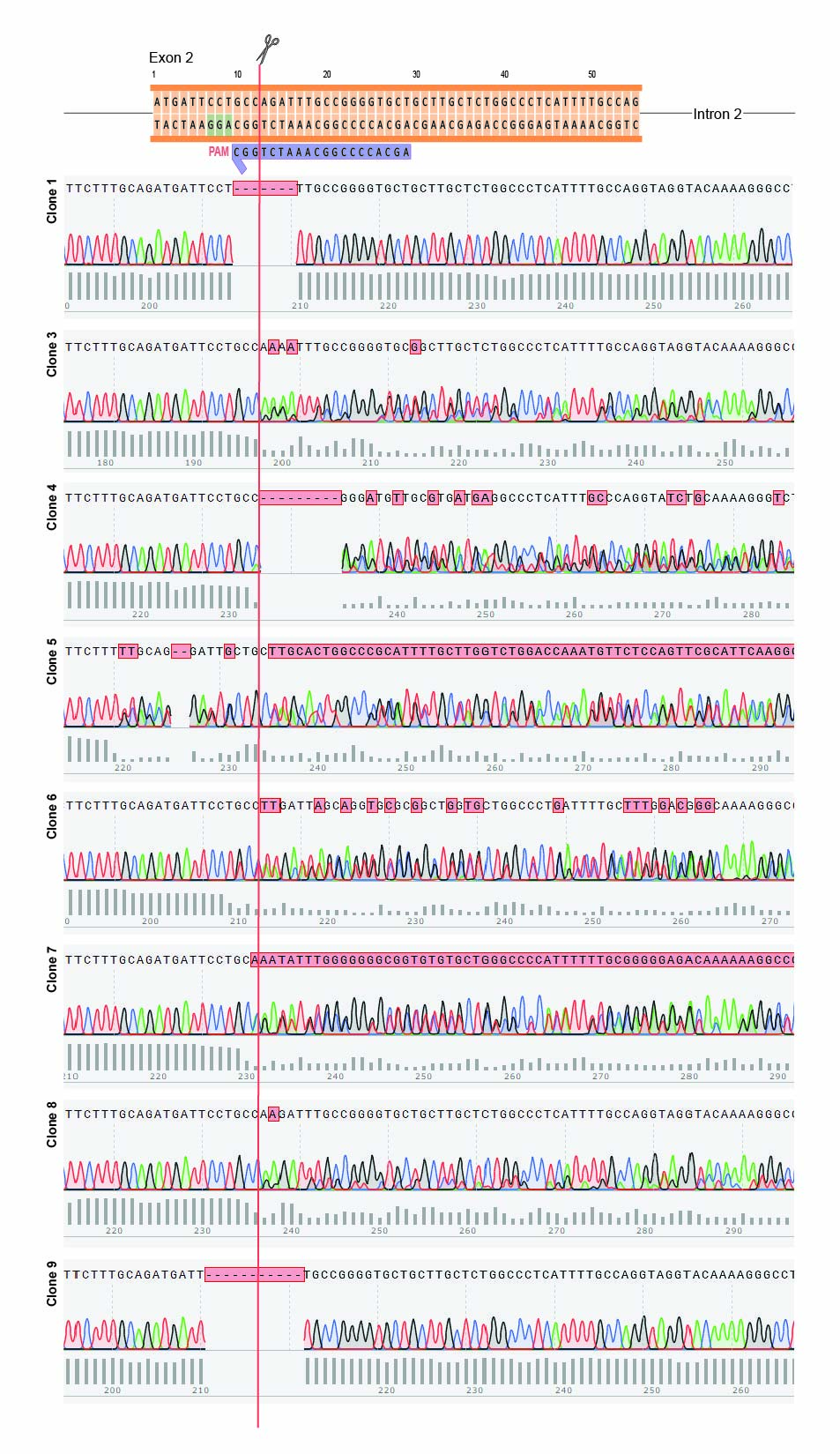
